# Exploiting the therapeutic vulnerability of IDH-mutant gliomas with zotiraciclib

**DOI:** 10.1101/2023.06.29.547143

**Authors:** Ying Pang, Qi Li, Zach Sergi, Guangyang Yu, Xueyu Sang, Olga Kim, Herui Wang, Alice Ranjan, Mythili Merchant, Belene Oudit, Robert W. Robey, Ferri Soheilian, Bao Tran, Felipe J. Núñez, Meili Zhang, Hua Song, Wei Zhang, Dionne Davis, Mark R. Gilbert, Michael M. Gottesman, Zhenggang Liu, Javed Khan, Craig J. Thomas, Maria G. Castro, Taranjit S. Gujral, Jing Wu

## Abstract

Isocitrate dehydrogenase (IDH)-mutant gliomas have distinctive metabolic and biological traits that may render them susceptible to targeted treatments. Here, by conducting a high-throughput drug screen, we pinpointed a specific susceptibility of IDH-mutant gliomas to zotiraciclib (ZTR). ZTR exhibited selective growth inhibition across multiple IDH-mutant glioma *in vitro* and *in vivo* models. Mechanistically, ZTR at low doses suppressed CDK9 and RNA Pol II phosphorylation in IDH-mutant cells, disrupting mitochondrial function and NAD+ production, causing oxidative stress. Integrated biochemical profiling of ZTR kinase targets and transcriptomics unveiled that ZTR-induced bioenergetic failure was linked to the suppression of PIM kinase activity. We posit that the combination of mitochondrial dysfunction and an inability to adapt to oxidative stress resulted in significant cell death upon ZTR treatment, ultimately increasing the therapeutic vulnerability of IDH-mutant gliomas. These findings prompted a clinical trial evaluating ZTR in IDH-mutant gliomas towards precision medicine (*NCT05588141*).

**Highlights:**

- Zotiraciclib (ZTR), a CDK9 inhibitor, hinders IDH-mutant glioma growth *in vitro* and *in vivo*.
- ZTR halts cell cycle, disrupts respiration, and induces oxidative stress in IDH-mutant cells.
- ZTR unexpectedly inhibits PIM kinases, impacting mitochondria and causing bioenergetic failure.
- These findings led to the clinical trial NCT 05588141, evaluating ZTR for IDH-mutant gliomas.

## Introduction

Treating diffuse gliomas poses an immense challenge due to the inter-tumor heterogeneity of the disease and the limited availability of effective drugs with sufficient penetration into the central nervous system (CNS). Level I evidence from clinical trial suggested the diverse clinical responses to the same treatment in subsets of histologically similar but molecularly distinct gliomas^1^. It highlights the significance of treating subsets of gliomas based on their unique molecular characterization, tumor biology and thereby therapeutic vulnerability, which will lead to a better clinical outcome.

Zotiraciclib (ZTR, TG02) is a pyrimidine-based multi-kinase inhibitor with inhibitory effects on the cyclin-dependent kinases (CDKs), primarily CDK9, and regulates transcriptional processes in multiple cancer cell models ^2^. The previous clinical trials have investigated ZTR and demonstrated its safety profiles ^3,4^. Our previous studies have shown that ZTR modulates various mechanisms of cancer cell survival in glioblastomas ^5^. ZTR suppressed both transcription and expression of antiapoptotic genes, leading to cell death in glioblastoma cells. Additionally, ZTR induced mitochondrial dysfunction and depleted cellular ATP. Importantly, *in vivo* experiments provided evidence to support the ability of ZTR to penetrate the blood-brain barrier (BBB), a prerequisite for targeting gliomas ^5^. Based on the promising preclinical and clinical evidence, the US Food and Drug Administration (FDA) granted orphan drug designation to ZTR as a therapy for malignant gliomas. Preliminary analysis of clinical outcomes revealed a statistically significant improvement in progression-free survival (PFS) among patients with IDH-mutant gliomas compared to those with IDH-wildtype gliomas. While a better PFS in patients with IDH-mutant gliomas may indicate a more favorable prognosis, other studies have suggested an accelerated disease progression after the initial recurrence, diminishing the prognostic value of an *IDH* mutation after the first disease progression ^6^. Most participants in this study were enrolled to the study after their second recurrence, suggesting a selective ZTR-related response in IDH-mutant gliomas.

*IDH* mutation is a crucial genetic feature found in a subset of adult diffuse gliomas and the mutation status is utilized as a key biomarker in the WHO classification of diffuse gliomas ^7^. Wildtype IDH converts isocitrate to a-ketoglutarate, generating NADPH, while the IDH-mutant enzyme produces the oncometabolite 2-hydroxyglutarate (2-HG) and consumes NADPH ^8,9^. Due to the accumulation of 2-HG and depletion of NADPH, IDH-mutant gliomas exhibit distinct biological characteristics, including redox imbalance, altered DNA damage repair, and heightened reliance on mitochondrial function for ATP production ^10–12^. These unique features of IDH-mutant gliomas create potential vulnerabilities to specific drugs that modulate tumor metabolism. Building upon the promising clinical activity of ZTR and a mechanistic rationale for targeting glioma vulnerability, we aimed to investigate the selective application of ZTR in gliomas harboring *IDH* mutations.

This study is designed to explore preclinical evidence for ZTR in IDH-mutant gliomas, unraveling the mechanism behind ZTR-induced mitochondrial dysfunction. Demonstrating ZTR’s selectivity and BBB penetration, which promise effective tumor control with reduced systemic toxicity, will advance precision neuro-oncology.

## Results

### Zotiraciclib has superior efficacy in IDH-mutant glioma cells

To demonstrate the superior efficacy of ZTR in IDH-mutant gliomas, we evaluated the cell viability of a panel of glioma cells with and without *IDH* mutations after ZTR treatment. Firstly, using patient-derived glioma stem-like cell models, we observed a lower half maximal inhibition concentration (IC_50_) of ZTR in IDH-mutant cells (TS603=7.06 nM and BT142=9.00 nM), compared to IDH-wildtype cells (GSC923=31.95 nM, GSC827=23.53 nM) (Figure 1A). In a pair of patient-derived adherent IDH-wildtype (U251) and IDH-mutant (HT1080) cells, HT1080 exhibited a lower IC_50_ of 35.94 nM compared to 66.13 nM in U251 when treated with ZTR (Figure 1B). Furthermore, we assessed the ZTR IC_50_ in mouse and patient-derived isogenic cell lines (NPA=30.73 nM/NPAI=17.65 nM, U87-wt=121.8 nM/U87-C8-mutant=71.6 nM /U87-C44-mutant=96.2 nM and GBM1-wt=23.13 nM/GBM1-mutant=16.95 nM). Consistently, IDH-mutant cells displayed a lower IC_50_ compared to the isogenic IDH-wildtype cells, indicating an increased sensitivity to ZTR in the IDH-mutant cells (Figure 1C). *IDH1* mutation status was confirmed in all cell lines (Suppl. Figure 1A). To further support the investigation of ZTR in IDH-mutant glioma, we conducted a high-throughput screen of 2,481 FDA-approved and investigational drugs using six primary patient-derived glioma cell lines with *IDH1* mutations ^13^. ZTR was found to be one of six drugs showing both high efficacy of cell killing and capacity to penetrate BBB (MPO>4 and AUC Z-score<-3), indicating ZTR is one of the most effective agents among all screened compounds that can be used in patients treating IDH-mutant gliomas (Figure 1D).

**Figure 1.**
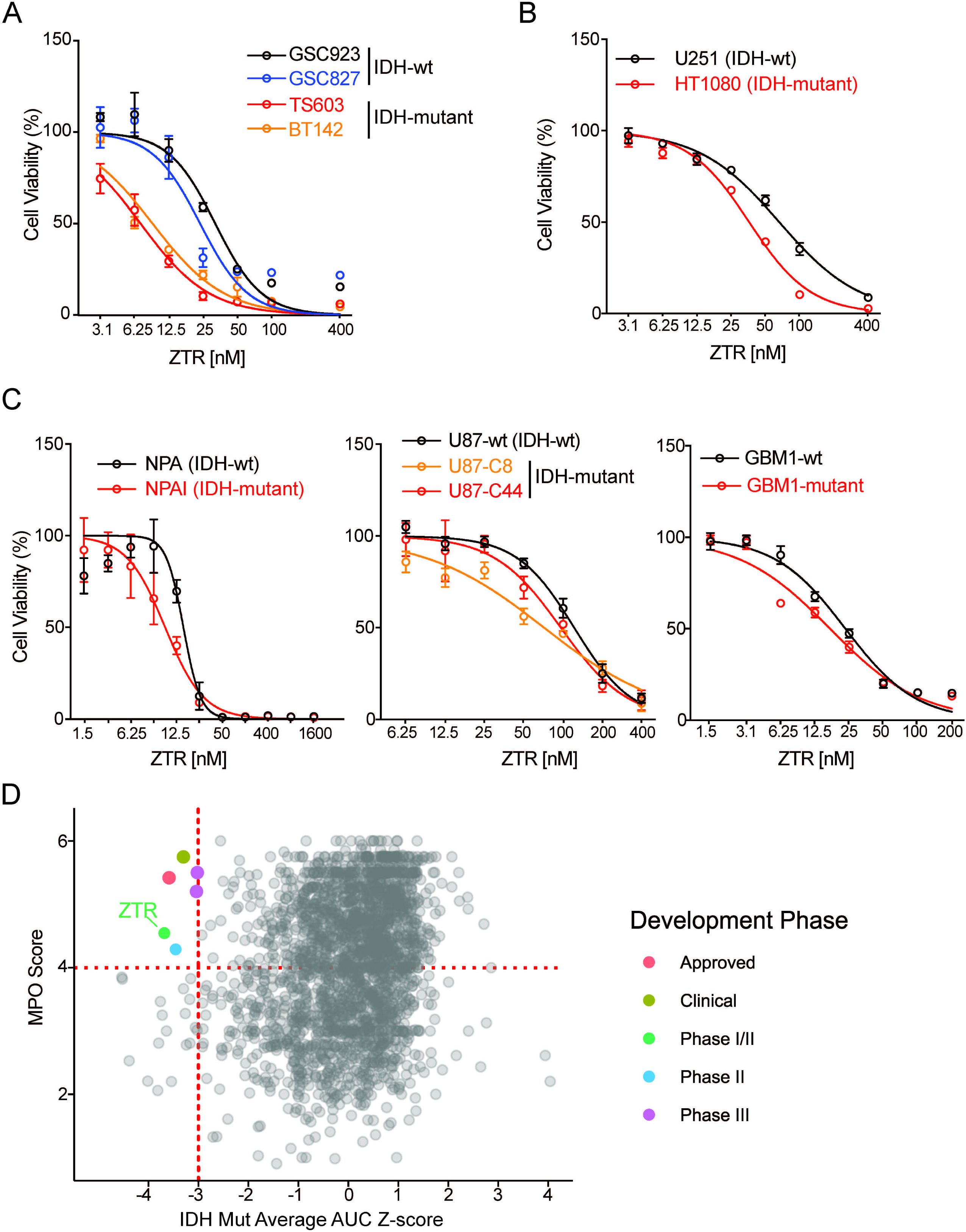
ZTR has enhanced anti-glioma effects on IDH-mutant glioma cells. A. The cytotoxic effect of ZTR in patient-derived stem-like glioma cells (a), patient-derived adherent tumor cells (b), isogenic mouse-derived cells (c), and isogenic human-derived cells (d-e). Cells were treated with serial dosages of ZTR for 72 hours prior to cell viability assay. B. Cell proliferation assay of patient-derived stem-like cells treated with 15 nM ZTR. Phase area confluence was measured every 3 hours by IncuCyte up to 72 hours. All cell line data were normalized to its own start time point (0 hour) value. C. In a high throughput drug screen of 2,481 FDA-approved or investigational drugs, the MPO score against average AUC Z-Score of six primary patient-derived glioma cell lines with *IDH1* mutations showing that ZTR (green dot) was one of the few agents having both high efficacy and capacity to penetrate BBB, defined as AUC Z-score<-3 and MPO>4, respectively.

### ZTR confers survival benefit in *in vivo* models of IDH-mutant gliomas

Following the observation of superior efficacy in IDH-mutant cells, *in vivo* experiments were conducted to determine the survival benefit of ZTR treatment. ATP-binding cassette (ABC) transporter proteins, including P-glycoprotein transporters (Pgp, encoded by *ABCB1*), multidrug resistance-associated protein (MRP1, encoded by *ABCC1*), and ABC family member G2 (encoded by *ABCG2*) are known to transport a wide range of chemotherapies and targeted drugs, leading to cancer drug resistance ^14^. To determine whether ZTR is a potential substrate of ABC transporter proteins, we tested the treatment response to ZTR in cells overexpressing each of these three major transporter proteins. Cells transfected with ABCB1(MDR-19 cells), ABCG2(R-5 cells), and ABCC1 did not show any difference in response to ZTR treatment compared to cells transfected with empty vector pcDNA (Figure 2A). However, these cells lines exhibited significant resistance to chemotherapeutic drugs known to be substrates of these transporters, such as mitoxantrone (Suppl. Figure 1B). These findings suggest that ZTR is unlikely to be a substrate for ABC transporter proteins, thereby a better chance to cross the blood-brain-barrier. Next, we evaluated the efficacy of ZTR in a syngeneic mouse model bearing orthotopic IDH-wildtype or IDH-mutant gliomas by intracranial implantation of NPA and NPAI cells, respectively. Seven days after the intracranial tumors were implanted, mice were treated with either vehicle or ZTR (Figure 2B). As expected, mice bearing IDH-mutant gliomas survived longer than those with IDH-wildtype tumors. ZTR treatment prolonged survival significantly from 30 to 33 days in IDH-mutant cohort (p=0.01) (Figure 2C). However, a statistically significant survival prolongation was not observed in ZTR-treated IDH-wildtype group (18 vs 20 days, p=0.07). Tumor samples were collected when mice were euthanized at the endpoints.

**Figure 2.**
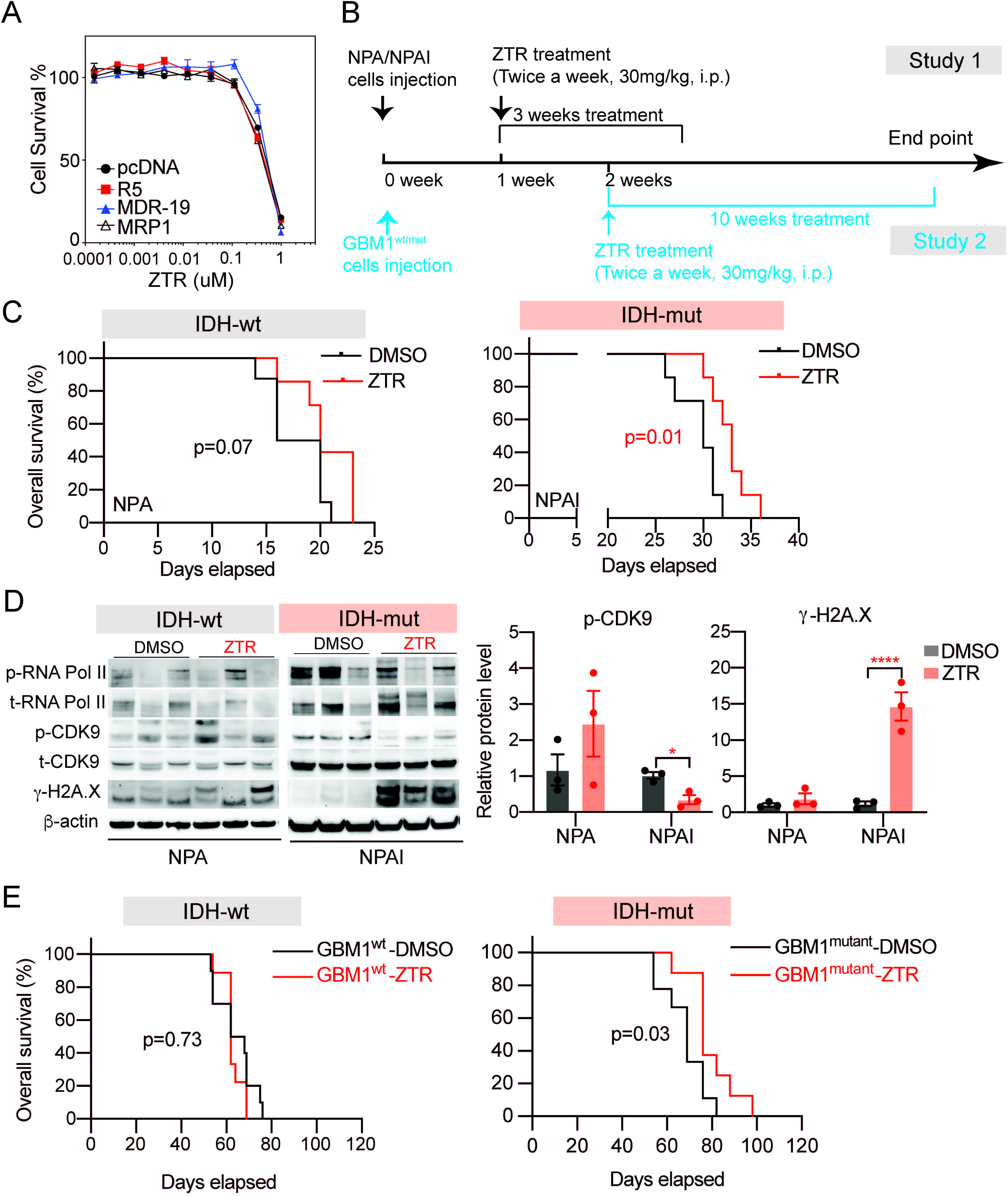
ZTR confers survival benefit in *in vivo* models of IDH-mutant gliomas. A. ABC transporter assay showing that overexpression of R5, MDR-19 and MRP1, the common ABC multidrug transporters, does not affect ZTR efficacy, suggesting ZTR is unlikely to be the substrate of these ABC transporters. Cells carrying pcDNA were used as negative controls and the expression of the transporters was confirmed by FACS analysis and demonstration that they conferred resistance to other cytotoxic agents that were substrates for the transporters. B. Schematic illustration for the allograft (experimental 1) and xenograft (experimental 2) modeling and ZTR treatment schedule. C. Kaplan-Meier analysis of orthotopic allograft in mice bearing NPA/NPAI cells that received ZTR treatment. D. (a) Western blotting showing protein expressions of CDK9, and RNA Pol II phosphorylation and γ-H2A.X from tumor tissues with or without ZTR treatment. GAPDH was blotted as internal control. (b) Statistic analysis of p-CDK9 and γ-H2A.X relative protein levels, which was normalized to b-actin. E. Kaplan-Meier analysis of intracranial xenograft in mice bearing GBM1-wt/-mutant cells that received ZTR treatment.

CDK9 phosphorylation was found to be suppressed in tumors from ZTR-treated mice compared to the DSMO-treated ones in the NPAI (IDH-mutant) group but not in the NPA (IDH-wildtype) mice. This finding of a brain pharmacodynamic marker affected by ZTR further supports the BBB penetration of the drug. The expression of γ-H2A.X, an indicator of DNA damage, was observed in both IDH-wildtype and IDH-mutant groups following treatment but was more significant in the IDH-mutant group (Figure 2D). The survival benefit of ZTR treatment in IDH-mutant tumors was further confirmed in a patient-derived xenograft murine glioma model (experimental 2) established by intracranial injection of isogenic GBM1-wildtype (65 vs 62 days, p=0.73) or GBM1-mutant (69 vs 76 days, p=0.03) cells (Figure 2B, E). Overall, ZTR prolonged the survival of mice bearing IDH-mutant gliomas. The findings of CDK9 activity suppression and DNA damage induction in tumor tissues from treated mice further support the BBB penetration of the drug and identified a potential pharmacodynamic biomarker of ZTR treatment for clinical investigation.

### The expression profiling indicates IDH-mutant cells struggle to thrive under the high stress levels induced by ZTR

To investigate the selective sensitivity of IDH-mutant vs IDH-wildtype cells to ZTR, we selected a lower dose (LD) of ZTR (15 nM) in contrast to 50 nM, which was used to treat patient-derived glioblastoma cells in our previous *in vitro* studies. Live-cell imaging analysis by IncuCyte revealed an enhanced suppression of cell proliferation in IDH-mutant cell lines (TS603 and BT142) following LD ZTR treatment, while no significant effect was observed in IDH-wildtype cells (GSC923 and GSC827) (Figure 3A). The proliferation data showed a significant difference from control after 42 hours in TS603 and 12 hours in BT142. To explore the critical gene and pathway alterations induced by the LD ZTR, we performed RNA sequencing (RNA-seq).

**Figure 3.**
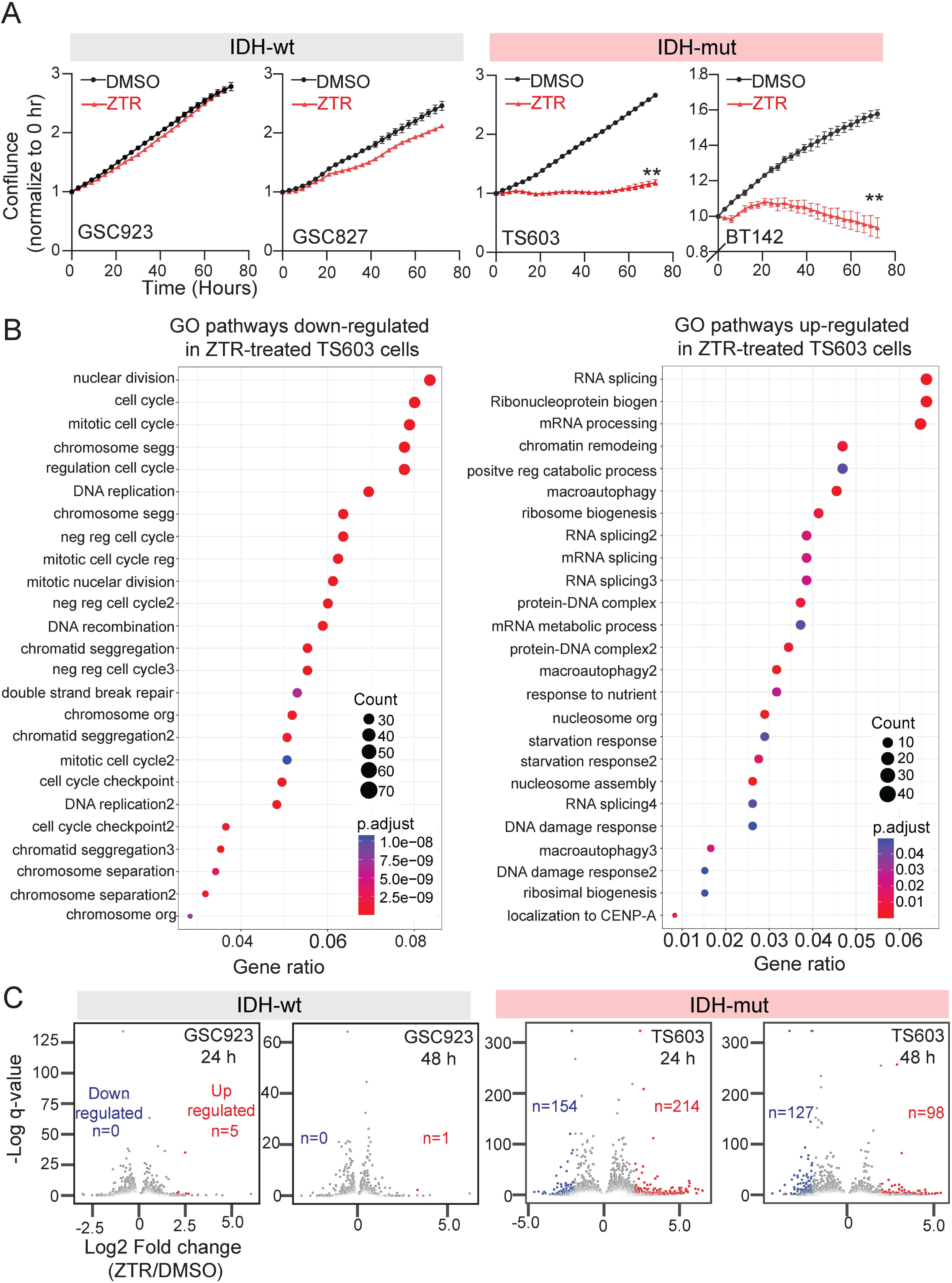
Low dose ZTR suppresses cell proliferation and affects stress response pathways in IDH-mutant glioma cells. A. Cell proliferation assay of patient-derived stem-like cells with IDH-wt and IDH-mut following 15 nM ZTR treatment. Phase area confluence was measured every 3 hours by IncuCyte up to 72 hours. All cell line data were normalized to its own start time point (0 hour) value. B. Enriched Gene Ontology (GO) pathways downregulated (left) or upregulated (right) in TG603 cells in response to ZTR treatment at 15 nM for 24 hours. C. Volcano plots based on RNAseq results showing the mRNA level changes of a panel of stress & toxicity pathway related, differentially expressed genes after ZTR treatment at 15 nM for 24 or 48 hours. n=3. ** p<0.01, Two-sample t-test at each time point with the Bonferroni correction.

Pathway analysis using Gene Ontology (GO) demonstrated several pathways down-regulated in ZTR-treated TS603 cells after 24 hours of treatment, indicating the cell cycle-related suppression. (Figure 3B). Moreover, the suppression of cell cycle-related pathways was maintained 48 hours after ZTR treatment (Suppl. Figure 1C). On the other hand, several pathways were identified to be up-regulated in the ZTR-treated TS603 cells, including ribosome biogenesis, mRNA processing and macroautophagy, suggesting the possibility of an adaptive response to cell proliferation/cell cycle suppression. We further examined the expression of genes related to stress & toxicity pathways. The volcano plot revealed a higher number of differentially expressed genes in TS603 compared to GSC923 upon LD ZTR treatment for 24 and 48 hours, indicating an elevated level of induced stress by LD ZTR treatment in IDH-mutant glioma (Figure 3C). The combination of a higher level of LD ZTR-induced stress and cell cycle suppression indicates that IDH-mutant cells fail to thrive in response to the treatment.

### ZTR suppresses mitochondrial complexes and induces mito-stress in IDH-mutant glioma cells

Mutant IDH is known to induce altered cell metabolism and increase dependence on oxidative mitochondrial metabolism ^15,16^. Previously, we demonstrated that ZTR induces mitochondria dysfunction in glioblastoma cells at 50nM ^17^. To investigate ZTR-induced mitochondrial changes in IDH-mutant glioma, we first examined the cellular organelle morphological changes induced by LD ZTR using a transmission electron microscopy (TEM) examination in GSC923 and TS603 cells. In LD ZTR-treated IDH-mutant TS603 cells, mitochondria appeared dysmorphic and decreased in size. There was an increased number of double-membraned vesicles, varying in size and filled with digested materials, suggesting autophagy (Figure 4A). We then focused on mitochondria and conducted an experiment using MitoTracker Green FM labeling. The results showed a greater loss of mitochondrial mass in the IDH-mutant glioma cells after LD ZTR treatment compared to the IDH-wildtype glioma cells (Figure 4B). Next, we examined the impact of LD ZTR on mitochondrial respiration complexes. The expression of genes encoding complexes I-V and other pathway activity signature genes (Suppl. Table 1) was suppressed by LD ZTR specifically in TS603 cells, but not in GSC923 cells (Figure 4C). This expression profile was also confirmed by the quantitative expression assay on pathway-focused genes encoding mitochondrial complexes using a profiler PCR array (Suppl. Figure 1D). Protein expression of most complexes was also decreased in IDH-mutant cells (TS603 and BT142) following LD ZTR treatment, as observed in native blue gel of the mitochondrial pellet (Figure 4D). To further confirm the suppression of complex activity, an in-gel activity assay for complex I was performed, revealing ZTR-induced activity suppression only in IDH-mutant TS603 and BT142 cells, but not in IDH-wildtype GSC923 and GSC827 cells (Figure 4E). Although there was a decrease in protein expression level after LD ZTR in GSC827 (Figure 4C), no decrease in activity was detected (Figure 4E). The suppressed activity of complexes led to a significant reduction in the oxygen consumption rate (OCR) in IDH-mutant cells, indicating decreased oxidative phosphorylation and increased oxidative stress. Surprisingly, the extracellular acidification rate (ECAR), which is expected to increase in stress responses, was reduced in IDH-mutant cells ^18^ (Figure 4F). The suppressive effect of glycolysis was also confirmed in the isogenic IDH-wildtype and -mutant models (Suppl. Figure 1E). Importantly, ATP production through mitochondrial respiration and glycolysis was significantly decreased in the IDH-mutant cells, suggesting a severe bioenergetic failure after LD ZTR treatment (Figure 4G, H).

**Figure 4.**
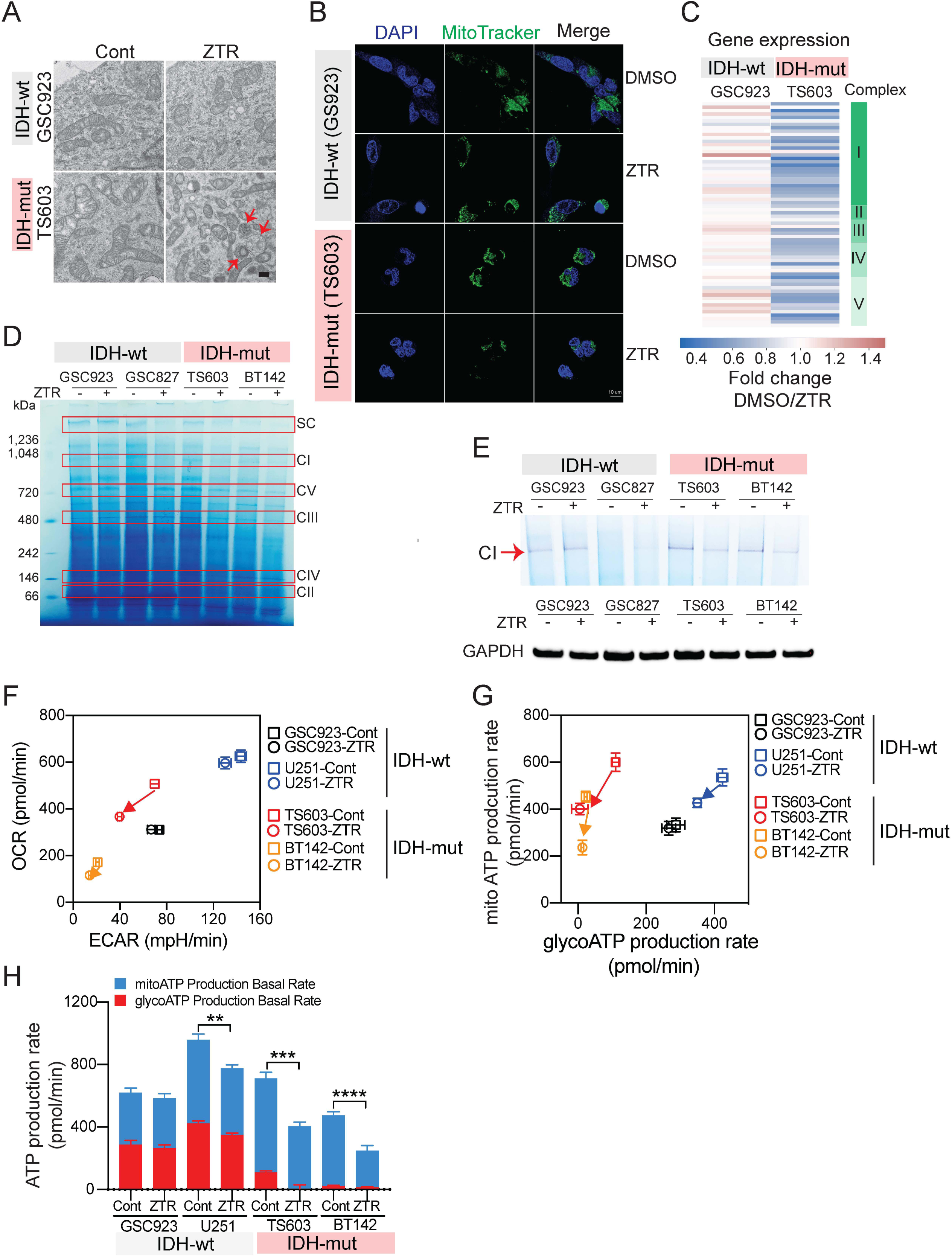
Low dose ZTR treatment suppresses mitochondrial complexes and induces biogenic energy depletion in IDH-mutant glioma cells. A. Transmission electron microscope pictures showing the mitochondrial morphology change and presence of double-membraned vesicles, filled with digested materials (indicated by red arrows), consistent with autophagy in TS603 cells after ZTR treatment at 15 nM for 48 hours. Scale bar=500 nm. B. MitoTracker Green FM staining demonstrating the mitochondrial mass volume change after ZTR treatment at 15 nM for 48 hours in IDH-wildtype and IDH-mutant cells. Scale bar=10 um. C. RNAseq results showing fold change in expression level of the signature genes coding for mitochondrial complexes after ZTR treatment at 15 nM for 48 hours, in GSC923 and TS603 cells. D. Native blue gel staining of mitochondrial complexes protein after ZTR treatment at 15 nM for 48 hours in IDH-wildtype (GSC923, GSC827) and IDH-mutant (TS603 and BT142) cells. E. In-gel activity assay showing the complex I activity changes after ZTR treatment at 15 nM for 48 hours. GAPDH was blotted as internal control. F. Mito-stress seahorse assay showing mitochondrial stress by measuring the OCR and ECAR after ZTR treatment at 15 nM for 24 hours. G. Real-time ATP seahorse assay measuring mitoATP and glycolATP production changes after ZTR treatment at 15 nM for 24 hours. *, *P* < 0.05; **, *P* < 0.01; ***, *P* < 0.001.

### ZTR-induced mito-stress intensifies the imbalance of redox pathway in IDH-mutant glioma cells

Complex I is essential for regenerating NAD in the mitochondria^19^. The deficiency in complex I induced by ZTR may further interrupt the NAD/NADH balance. In our IDH-mutant cell model, we observed a lower baseline level of NAD expression compared to the IDH-wildtype cell (Figure 5A and Suppl. Figure 1F), which is consistent with the previously reported findings^20^. While ZTR-induced NAD suppression was observed in both IDH-mutant and IDH-wildtype cells, the decrease in NAD levels was much more pronounced in IDH-mutant gliomas upon ZTR treatment (Figure 5A). Nicotinamide phosphoribosyltransferase (NAMPT) and nicotinate phosphoribosyltransferase domain containing 1 (NAPRT1) are two major rate-limiting enzymes involved in NAD synthesis that determine NAD levels^21^. The NAMPT activity assay failed to show a ZTR-induced NAMPT activity suppression up to a concentration of 50 nM, while a positive control, FK866, significantly reduced NAMPT activity (Figure 5B). Protein expression of NAMPT and NAPRT1 was not affected by LD ZTR treatment, suggesting that the reduction in NAD induced by ZTR is unlikely due to the suppression of NAMPT and NAPRT1. However, we observed a significant reduction in expression of poly (ADP-ribose) glycohydrolase (PARG) following LD ZTR in IDH-mutant glioma cells (Figure 5C). PARG is an enzyme involved in degrading ADP-ribose polymers to release NAD for re-use. The mechanism underlying the reduction in PARG is unclear, but it is hypothesized to contribute to the further reduction in NAD levels. In summary, LD ZTR reduced NAD levels in IDH-mutant glioma cells through the suppression of complex I and PARG expression. The impact of ZTR-induced NAD reduction was expected to be greater in IDH-mutant gliomas, which inherently have lower levels of NAD compared to their IDH-wildtype counterpart.

**Figure 5.**
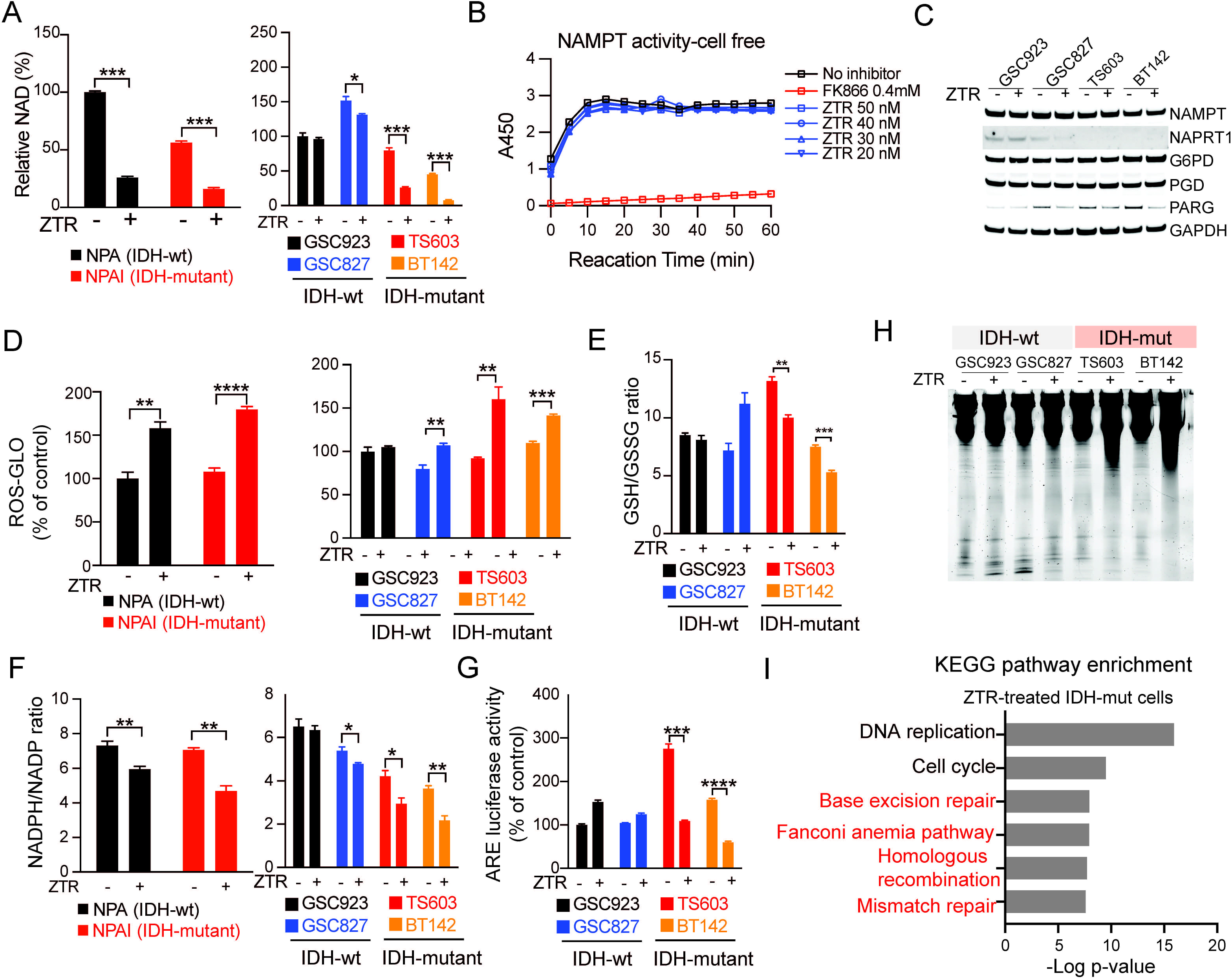
Low-dose ZTR-induced mito-stress intensifies the imbalance of redox pathway in IDH-mutant glioma cells. A. NAD-Glo assay measuring NAD level changes in NPA/I or GSC cells after ZTR treatment at 50 nM and 15 nM, respectively. The signals were measured after 48 hours’ treatment. B. Cell-free NAMPT activity assay measuring NAMPT enzyme activity with different dose of ZTR treatment in time-dependent manner. FK866 was used as a positive control. C. Western blotting shows protein levels of NAD and NADPH related enzymes with and without ZTR treatment at 15 nM for 72 hours. GAPDH was blotted as internal control. D. ROS-Glo assay measuring the ROS level in NPA/I or GSCs cell lines with and without ZTR treatment at 50 nM and 15 nM, respectively. E. GSH/GSSG-Glo assay measuring the GSH/GSSG ratio changes in patient-derived GSCs cell lines following ZTR treatment at 15 nM. F. NADP/NADPH-Glo assay measuring NADP/NADPH changes in NPA/I or patient-derived GSC cells after ZTR treatment at 50 nM and 15 nM, respectively. G. ARE luciferase assay measuring the ARE activity changes in GSCs cell lines with ZTR treatment at 15 nM. H. Total genomic DNA electrophoresis analysis showing DNA fragmentation after ZTR treatment at 15 nM for 48 hours. I. RNAseq results of GSC923 and TS603 cell lines showing significantly enriched gene sets related to DNA damage and repair processes after 24 hours ZTR treatment at 15 nM. *, *P* < 0.05; **, *P* < 0.01; ***, *P* < 0.001.

Oxidative stress induces the production of mitochondrial reactive oxygen species (ROS) and disrupts redox homeostasis ^22^. LD ZTR significantly increased ROS production in the IDH-mutant cells, indicating mito-stress is induced after LD ZTR treatment (Figure 5D). To counterbalance the excessive ROS, a substantial amount of antioxidant is consumed, as evidenced by a significant decrease in GSH/GSSG ratio in IDH-mutant cells (Figure 5E). NADPH, which is critical for glutathione and thioredoxin systems to neutralize ROS, is largely consumed by 2-HG production in IDH-mutant cells ^23^ (Suppl. Figure 1F). LD ZTR treatment further reduced NADPH/NADP ratios to a greater extent in IDH-mutant cells compared to IDH-wildtype cells. In patient derived IDH-mutant GSCs, the NADPH/NADP ratio was downregulated by 30.3% and 40.5% in TS603 and BT142, respectively. However, there was only a 11.3% decrease in the IDH-wildtype GSC827 and no change in IDH-wildtype GSC923 (Figure 5F). The crucial enzymes G6PD and PGD, responsible for NADPH production, were not altered after ZTR treatment (Figure 5C). These results suggest that the low NADPH is likely caused by the increased consumption rather than decreased production of NADPH, thereby exacerbating the ROS imbalance in IDH-mutant gliomas.

As a cellular self-defense mechanism in response to increased ROS production, a group of genes encoding antioxidants is expected to be activated to eliminate excess ROS. However, the activity of antioxidant response element (ARE) luciferase, which reflects transcriptional activation of those genes, was found to be decreased rather than increased in IDH-mutant cells (Figure 5G). The mechanisms underlying this paradoxical reduction in ARE luciferase activity are unclear, but it further contributes to the redox imbalance. Excessive ROS production is a well-recognized mediator of induced DNA damage. In IDH-mutant cells, more genomic DNA fragmentation was observed compared to IDH-wildtype cells (Figure 5H). We further analyzed the enrichment of signaling pathways based on the RNA-seq data from GSC923 and TS603. KEGG pathway enrichment analysis showed ZTR induced suppression in multiple DNA damage repair pathways (Figure 5I). The ZTR-induced depletion of NAD, which serves as the substrate for the PARP signaling pathway, might contribute to the defect of DNA damage repair function, particularly in base excision repair, in IDH-mutant cells.

### LD ZTR induces cell death in IDH-mutant glioma cells

We previously demonstrated the suppression of RNA-Pol II and CDK9 activity in glioblastoma cells treated with ZTR at a concentration of 50 nM ^5^. In this study, we observed a similar suppression in IDH-mutant, but not IDH-wildtype glioma cells, when treated with ZTR with a lower dose at 15 nM (Figure 6A). The expression of short-lived anti-apoptotic proteins, namely MCL-1, XIAP and survivin, were also suppressed by LD ZTR in IDH-mutant cells (Figure 6Ba). Activation on c-PARP and γ-H2A.X further indicate the induction of DNA damage by LD ZTR in IDH-mutant cells, but not in IDH-wildtype cells (Figure 6Bb).

**Figure 6.**
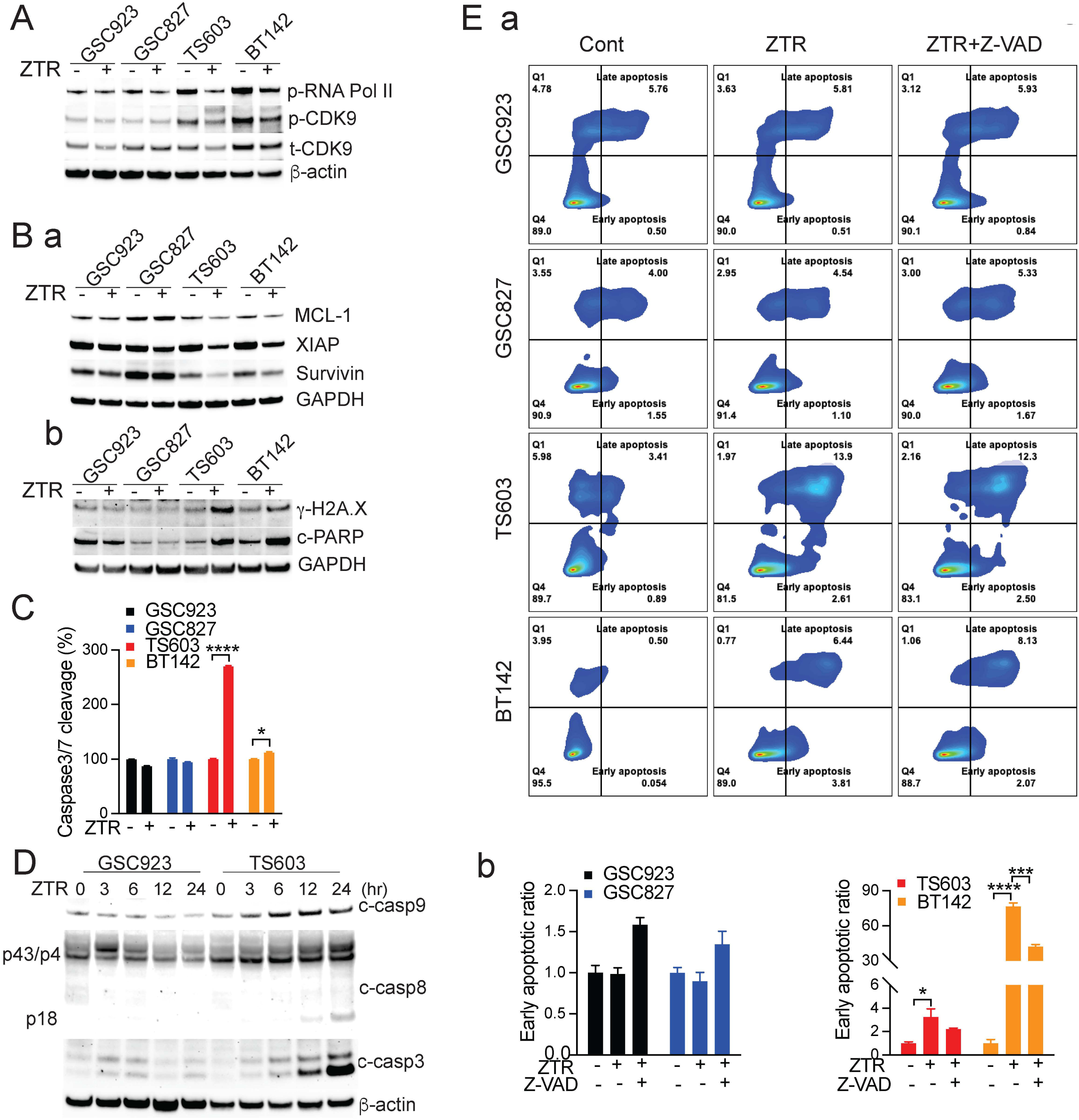
Low dose ZTR induces cell death in IDH-mutant glioma cells. A. Western blotting showing CDK9 and RNA Pol II phosphorylation levels with ZTR treatment at 15 nM for 72 hours. β-actin was blotted as internal control. B. (a) Western blotting showing short-lived antiapoptotic protein expression levels with ZTR treatment at 15 nM for 48 hours. (b) Western blotting showing DNA damage protein expression levels with ZTR treatment at 15 nM for 72 hours. GAPDH was blotted as internal control. C. Caspase 3/7-Glo assay showing caspase 3/7 activity changes after ZTR treatment at 15 nM for 72 hours. D. Western blotting showing cleavage of caspase 9, 8 and 3 after ZTR treatment at 15 nM. Samples were collected after 0, 3-, 6-, 12-, and 24-hours treatment. β-actin was blotted as internal control. E. (a) Flow cytometry assay measuring the percentage of early/late apoptotic and necrotic cell by staining with APC Annexin V/PI after 15 nM ZTR or both 15 nM ZTR and 10 uM Z-VAD for 72 hours. DMSO treatment was used as the control. (b) Statistic analysis of early apoptotic ratio in all cell lines.

Caspase 3/7 activity, a biomarker of apoptosis, was significantly increased in IDH-mutant cells following LD ZTR treatment, while no significant change was observed in IDH-wildtype cells (Figure 6C). The activation of caspase 8 and caspase 9 suggested that both intrinsic and extrinsic apoptosis pathways may have been activated as early as 6-12 hours in LD ZTR-treated IDH-mutant cells (Figure 6D). Apoptosis was further analyzed by flow cytometry, which showed an increase in the percentage of both early and late apoptotic cells in LD ZTR-treated IDH-mutant cells, but not in IDH-wildtype cells (Figure 6Ea). Interestingly, a pan-apoptosis inhibitor, Z-VAD, only partially rescued the cell death (Figure 6Eb). These results suggest that apoptosis, a programed ATP-dependent cell death, may not be major type of cell death in ZTR-treated cells. The cellular organelle morphological changes after the LD ZTR treatment suggested that autophagy may have been induced (Figure 4A). In addition, the RNAseq analysis in LD ZTR-treated cells indicated the up-regulated autophagy pathway (Figure 3C). LD ZTR treatment-induced cleavage of LC3B, an autophagy marker was demonstrated in IDH-mutant glioma cells (Suppl Figure 2B). Overall, multiple cell death mechanisms may be involved in LD ZTR-induced cell death in IDH-mutant cells.

### PIM is identified as a target of ZTR in glioma

Our data reveals that ZTR induces cell cycle arrest, disrupts cellular respiration, and triggers oxidative stress in IDH-mutant cells (Figure 4, 5, 6). However, the precise molecular mechanism by which ZTR induces mitochondrial dysfunction remains unresolved. To elucidate the specific kinases that may be involved in ZTR-mediated bioenergetic failure, we analyzed gene expression changes in TS603 cells treated with ZTR for 24 hours (Figure 7A). Pathway enrichment analysis was performed on 915 significantly downregulated genes by ZTR, with at least a 2-fold decrease in expression. We identified an enrichment of genes related to Mitochondrial complex I assembly, with a p-value of 0.04. Subsequently, we utilized the ZTR-downregulated genes associated with mitochondrial complex assembly to infer potential upstream kinases ^24^ that might regulate these genes. Utilizing the eXpression2Kinases (X2K) method for kinase enrichment analysis identified the Provirus Integration site for Moloney leukemia virus (PIM) family of kinases as potential candidates capable of regulating ZTR-induced mitochondrial dysfunction.

**Figure 7.**
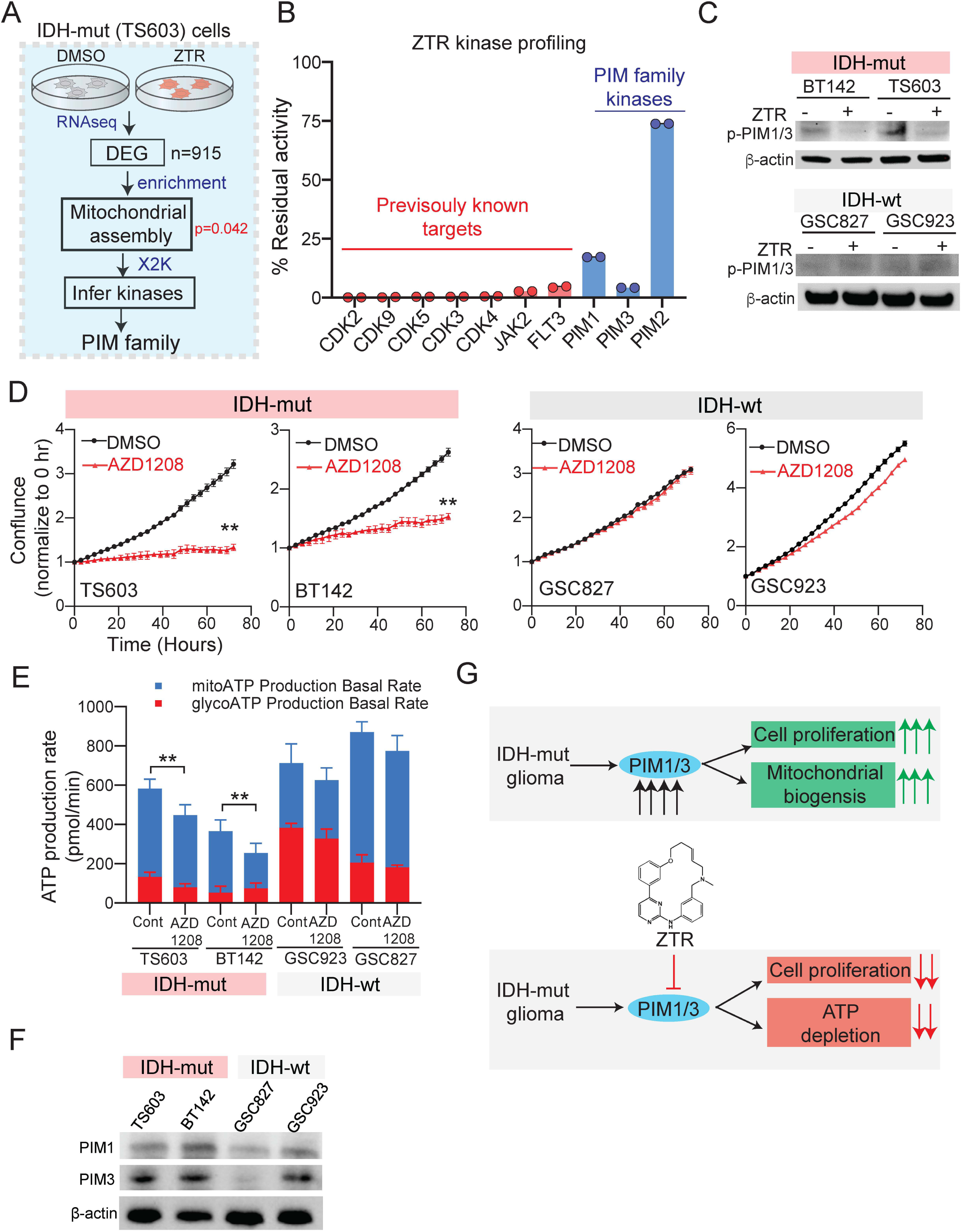
PIM1/3 are targets of ZTR, mediating enhanced effects in IDH-mutant gliomas. A. A schematic illustrating an integrated pathway enrichment and upstream kinase inference approach was utilized to identify PIM kinases as potential targets of ZTR associated with mediating mitochondrial dysfunction. B. ZTR (500nM) inhibits the activity of several previously known CDK kinases, and PIM kinases *in vitro*. Bars represent the mean of two independent data points. C. Western blot showing the activity changes of PIM1/3 following treatment of ZTR in IDH-mutant and IDH-wildtype glioma cells. D. AZD1208, a pan inhibitor of PIM significantly suppresses the cell proliferation in IDH-mutant cells but not in IDH-wildtype cells, demonstrated by the cell proliferation assay using IncuCyte. Phase area confluence was measured every 3 hours by IncuCyte up to 72 hours. All cell line data were normalized to its own start time point (0 hour) value. E. Real-time ATP seahorse assay measuring mitoATP and glycolATP production changes after treatment of AZD1208, a PIM inhibitor for 24 hours. F. Western blot showing the increased protein expression of PIM1/3 in IDH-mutant compared to IDH-wildtype glioma cells. G. A graphic summary to illustrate that ZTR targets PIM1/3 to induce cell proliferation suppression and cellular ATP depletion. * *p*<0.05, \*\**p*<0.01

To validate this hypothesis, we performed kinase activity assays to determine whether ZTR directly inhibits PIM kinases. Interestingly, we found that ZTR is a potent inhibitor not only of several CDK kinases, JAK2, and FLT3, as demonstrated previously ^2^, but it also effectively inhibits PIM1 and PIM3 kinases (Figure 7B). Consistently, phosphorylation of PIM1/3 was notably reduced following LD ZTR treatment in IDH-mutant glioma cells but not in IDH-wildtype ones. These results imply that low-dose ZTR exerts a greater inhibitory influence on PIM1/3 activity specifically in IDH-mutant cells. (Figure 7C). To explore the biological effect of PIM1/3 inhibition in glioma cells, we performed the live-cell imaging analysis by IncuCyte following the treatment with AZD1208, a pan-PIM inhibitor. PIM inhibition induced a significant suppression of cell proliferation in IDH-mutant cell lines (TS603 and BT142), but not in IDH-wildtype cells (Figure 7D). Treatment with PIM inhibitor induced a significant reduction in ATP production, specifically in ATP generated from oxidative phosphorylation, in IDH-mutant cells rather than IDH-wildtype cells (Figure 7E). The effects of inhibiting PIM on cell proliferation and ATP reduction are consistent with those caused by LD ZTR treatment (Figure 3A and Figure 4H), implying that PIM is a viable target for the inhibitory action of LD ZTR. Notably, PIM exhibits constitutive activation. Consequently, the higher protein expression levels of PIM1/3 in IDH-mutant cells compared to IDH-wildtype cells (Figure 7F) indicate elevated PIM activity in the mutant cells. In conclusion, as depicted in Figure 7G, our findings suggest that ZTR’s impact on mitochondrial complex I assembly and bioenergetic failure in IDH-mutant cells may be mediated, in part, by its potent and unexpected inhibition of PIM1 and PIM3 kinases.

## Discussion

In this study, we have provided preclinical evidence and the underlying mechanisms for the therapeutic vulnerability to ZTR treatment in IDH-mutant gliomas. We observed selective growth inhibition across multiple *in vitro* models of IDH-mutant gliomas and a survival benefit in both syngeneic and PDX mouse models with orthotopic IDH-mutant gliomas. To further support further clinical investigation of ZTR in IDH-mutant glioma patients, we identified that ZTR is one of the few among a panel of 2481 FDA-approved or investigational drugs, exhibiting selective sensitivity in IDH-mutant glioma while penetrating BBB in a high throughput drug screen. Mechanistically, we revealed that LD ZTR specifically suppressed mitochondrial complex activities in IDH-mutant gliomas, leading to a decreased NAD level, impaired ATP production, and thereby inducing mito-stress. Notably, we identified PIM kinases as a new target of ZTR, linking the ZTR-induced bioenergetic failure and cell growth suppression with PIM inhibition in glioma. The inherent redox imbalance of IDH-mutant gliomas and the compromised ROS clearance capability further contributed to the accumulation of ROS upon ZTR treatment. Increased ROS and suppressed DNA damage repair led to enhanced DNA damage. Furthermore, ZTR suppressed the CDK9/RNA POLII phosphorylation pathway, resulting in decreased expression of antiapoptotic proteins and contribute to cell death. In summary, as illustrated in graphic abstract, ZTR induced bioenergetic failure, DNA damage, and apoptosis, ultimately leading to the death of IDH-mutant glioma cells.

### Connecting ZTR-induced mitochondrial dysfunction with PIM Kinases

PIM kinases are in the family of serine/threonine protein kinases. Three isoforms, PIM1, 2 and 3 have been identified, where PIM 1 and 2 share a very similar kinase domain ^25^. All 3 isoforms of PIM are constitutively active, phosphorylate serine/threonine amino acids and activate similar cellular pathways ^26,27^. PIM kinases have a long list of phosphorylation targets, involving multiple signaling pathways, playing a role in tumorigenesis and tumor progression in both solid and hematologic cancers ^27,28^. More importantly, PIM inhibition has been reported to cause mitochondrial fragmentation and increased intracellular ROS in lung cancer ^29^. In our study, we identified PIM as a target of ZTR through an integrated biochemical profiling of ZTR targets and transcriptomic analysis. We also validated the expression of PIM1/3 in our glioma cell line models. More notably, we linked the ZTR-induced cell proliferation suppression and mitochondrial dysfunction in gliomas with PIM inhibition. To our knowledge, PIM has never been reported as a target of ZTR in glioma. Combined with the elevated protein expression in IDH-mutant cells, the findings provide a profound mechanistic understanding of how LD ZTR triggers mitochondrial dysfunction in IDH-mutant gliomas.

### The impact of ZTR-induced mitochondrial stress on IDH-mutant glioma cells can be attributed to their unique metabolic vulnerability

Mutant IDH leads to the accumulation of 2-HG and rewiring the citric acid cycle metabolism, resulting in the enhanced reliance of IDH-mutant cells on mitochondrial oxidative phosphorylation for cellular energy production ^10,15^. While increased mitochondrial density and activity was reported in IDH-mutant oligodendroglioma cells, indicating heightened mitochondrial metabolism in these cells ^30^, accumulated 2-HG in IDH-mutant gliomas was found to cause cell hypersuccinylation, leading to mitochondrial dysfunction ^31^. The combination of increased dependence on mitochondrial metabolism and underlying mitochondrial dysfunction in IDH-mutant gliomas may explain their vulnerability to ZTR treatment, which further suppress the mitochondrial function ^5,32^. In our study, we observed that LD ZTR suppressed the expression of respiration complexes at both RNA and protein levels, inducing mito-stress. This treatment also resulted in an overall decrease in mitochondria mass and dysmorphic mitochondria in IDH-mutant glioma cells, disrupting oxidative phosphorylation and ATP production ^18,33,34^.

In addition to the reliance on mitochondria metabolism, IDH-mutant gliomas also exhibit a reduction in NAD levels due to the epigenetic downregulation of nicotinate phosphor-ribosyltransferase (Naprt1), an enzyme involved in NAD+ salvage pathway ^20^. In our study, LD ZTR treatment further suppressed NAD+ levels in the IDH-mutant cells, which may contribute to impaired NAD+-mediated PARP DNA repair pathway. Rather than affecting NAMPT and NAPRT1, the rate-limiting enzymes of the NAD salvage pathway, ZTR was found to suppress the expression of PARG. PARG is responsible for the degradation of PAR chain and the removal of ADP-ribose moieties from modified proteins, thereby recycling NAD+ back into the system ^35^. LD ZTR-induced PARG reduction may further contribute to the shortage of NAD in IDH-mutant glioma cells. Additionally, the mitochondrial complex I catalyzed the NADH oxidation and the elevation of NAD+, which is essential for energy production. The suppression of complex I activity by the LD ZTR treatment may partially contribute to the reduction of NAD induced by ZTR.

NADPH, another important molecule in the antioxidant system, plays a critical role in clearing up accumulated ROS. Depletion of NADPH results in the failure to effectively eliminate ROS, thereby potentiating oxidative stress ^36^. In IDH-mutant glioma cells, the conversion of α-KG to 2-HG excessively consumes NADPH, resulting in an imbalance of redox homeostasis ^37,38^. LD ZTR treatment, by suppressing electron transport chain complex, particularly complex I, further increases ROS production and exacerbates the pre-existing redox imbalance, resulting in a more pronounced impact on IDH-mutant glioma cells.

### ZTR suppressed several stress response pathways in IDH-mutant glioma cells

When cells are under mito-stress, glycolysis is expected to be upregulated to compensate for the loss of cellular energy^20^. However, in our study, a glycolysis suppression rather than enhancement was observed while oxidative phosphorylation was reduced in IDH-mutant cells following LD ZTR. This suppression in glycolysis may be attributed to the depletion of NAD depletion or the inhibition of glycolytic enzymes by ROS ^35,36^. The simultaneous suppression of both oxidative phosphorylation and glycolysis by ZTR resulted in a severe bioenergetic failure in IDH-mutant gliomas. This disruption of energy metabolism further contributes to the vulnerability of IDH-mutant gliomas to ZTR-induced cytotoxicity.

In response to the increased ROS production, cells typically activate the ROS scavenger systems, such as the GSH/GSSG system and NADPH /NADP to restore redox balance. Additionally, ARE plays a crucial role in mediating transcriptional activation of genes involved in detoxification and cytoprotection under oxidative stress ^39^. However, in our study, despite the presence of oxidative stress, LD ZTR treatment failed to activate ARE in IDH-mutant glioma cells. The reasons underlying this suppression are not yet clear, but it suggests a failure to restore redox balance following LD ZTR treatment. Consequently, the excessive accumulation of ROS resulted in multiple oxidative cellular damage, including DNA damage ^40^. Moreover, IDH-mutant tumors are characterized by global DNA hypermethylation, which can lead to the downregulation of the DNA damage sensor and impaired DNA repair pathway ^41^. This inherent DNA damage repair deficiency in IDH-mutant cells may further sensitize them to agents that induce DNA damage, such as ZTR. Consistent with this, our study observed the activation of DNA damage markers, γ-H2A.X and cleaved-PARP, and DNA fragmentation in LD ZTR-treated IDH-mutant cells.

The presence of *IDH* mutation in glioma cells has been shown to dynamically reprogram the transcriptome, leading to persistent downregulation of certain transcription factors ^42^. Additionally, the widespread hypermethylation, particularly in the promoter regions and CpG islands close to the transcription start sites, occurs in IDH-mutant cells, resulting in the predominant suppression of gene transcription ^43^. Therefore, the aberrant transcriptome of IDH-mutant tumors might further contribute to the vulnerability to ZTR-induced CDK9 and RNA POLII inhibition. We previously demonstrated that treatment with ZTR at a concentration of 50 nM led to a decrease in CDK9 and RNA POLII phosphorylation in IDH-wildtype GSC827 and GSC923 cells. However, when the dosage of ZTR was decreased to 15 nM, this suppression of transcription signaling was only observed in IDH-mutant cells and no longer detected in IDH-wildtype glioma cells.

Mutant IDH and the accumulation of 2-HG have been reported to induce apoptosis and autophagy in glioma cells, making them more susceptible to inhibition of the anti-apoptotic protein, Bcl-xl ^44,45^. In addition to this vulnerability, LD ZTR further decreased the expression of MCL-1, XIAP, and survivin in IDH-mutant glioma cells, while the same suppression was not observed in IDH-wildtype gliomas. Although apoptosis and autophagy were observed in the cell death process, the inhibition of these processes alone was not sufficient to completely prevent cell death, indicating the involvement of other cell death mechanisms. While markers of pyroptosis and necroptosis were not detected in ZTR-treated glioma cells, including those with *IDH* mutations, other cell death mechanisms, such as ferroptosis-induced cell death, may be worth investigating.

### Preclinical findings provided rationale for clinical investigation of ZTR in IDH-mutant gliomas

Indeed, the multiple mechanisms of action of ZTR contribute to its anti-glioma effects, targeting various biological features of IDH-mutant gliomas. The treatment-induced mitochondrial dysfunction and transcriptional regulation target the inherent biological features of IDH-mutant gliomas, including increased dependence on oxidative phosphorylation, redox imbalance, and NAD reduction. The increased vulnerability to the specific mechanisms of action of ZTR has led to increased efficacy of the drug treatment in this subset of gliomas with *IDH* mutation. The selective sensitivity of IDH-mutant gliomas to ZTR presents an opportunity to treat IDH-mutant glioma patients with less intense dosing schedule, thereby reducing the drug toxicity while treating the tumor and improving the patients’ quality of life.

Acknowledging the limitations of the study is crucial for the interpretation of the results and the development of future research. One major limitation is the modest survival benefit observed in IDH-mutant glioma mouse models when ZTR was administered as a single agent. Although the survival prolongation was statistically significant, the improvement was considered modest. It is important to consider that the mouse models used in the study have a more aggressive disease course which limits the length of treatment. In a clinical setting, IDH-mutant gliomas generally have a less aggressive disease course than their IDH-wildtype counterparts, allowing for a longer therapeutic window and potentially greater survival benefits. Additionally, considering combination therapies, such as combining ZTR with radiation, may enhance therapeutic efficacy and improve patient outcomes. To address these limitations and further explore the potential of ZTR as a treatment option, ongoing clinical trials have been initiated. These clinical studies will provide valuable data on the efficacy and safety of ZTR in patients with IDH-mutant gliomas.

In conclusion, this study has shed light on the therapeutic potential of ZTR in targeting IDH-mutant gliomas. We first demonstrated the findings that highlight the ability of ZTR to disrupt mitochondrial function, particularly via PIM inhibition, suppress stress response pathways, and induce cell death specifically in IDH-mutant gliomas with a low concentration. Based on these promising results, a clinical study has been initiated to evaluate the efficacy and safety of ZTR as a monotherapy in IDH-mutant glioma patients (*NCT 05588141*). By exploiting the inherent vulnerabilities of IDH-mutant gliomas and leveraging the unique mechanisms of actions of ZTR, this therapeutic approach holds potential to provide effective tumor control while minimizing treatment toxicities.

## Author contributions

Conceptualization, J.W., Y.P.; methodology, J.W., Y.P., Q.L., G.Y., Z.S., F.J.N., M.M.G., M.G.C., Z.L., J.K., C.J.T., and T.G.; investigation, J.W., Y.P., Q.L.,G.Y., Z.S., Q.L., X.S., H.W., O.K., A.R., M.M., B.O., R.W.R., F.S., B.T., M.Z., H.S. W.Z., D.D. and T.G.;; statistical analyses, J.W., Y.P., Z.S. and T.G.; writing – original draft, J.W. and Y.P., writing – review & editing, J.W., Y.P., Q.L., Z.S., O.K., R.W.R., M.R.G., M.M.G., M.G.C., J.K., C.J.T., and T.G.; funding acquisition, J.W.; supervision, J.W.

All authors read and approved the final article and take responsibility for its content.

## Data Availability

Data will be made available upon reasonable request.

---

Supplementary Figure 1. A. Confirmation of the presence of IDH-mutant is demonstrated by either a) Western blotting, or b) Sanger sequencing in all human- and mouse-derived cell lines used in this study. B. ABC transporter assay showing that cells with overexpressed R5, MDR-19 and MRP1, the common ABC multidrug transporters, were resistant to mitoxantrone, which was a known substrate of these transporters. C. RNAseq results of GSC923 and TS603 cell lines showing enriched gene sets related to cell cycle processes after ZTR treatment at 15 nM for 48 hours. D. Profiler PCR arrays showing the changes in mRNA expression level of the signature genes coding for mitochondrial complexes and signature pathways following 48 hours treatment of ZTR at 15nM in GSC923 and TS603 cells. E. Mito-stress seahorse assay showing ECAR measurement after ZTR treatment at 50 nM for 24 hours in NPA and NPAI cells. F. Baseline levels of NAD and NADPH/NADP in NAP, NAPI and patient derived GSC cell lines.

Supplementary Figure 2. A. Western blot showing GSDMC, GSDMC and p-MLKL expression in all cell lines with ZTR treatment.

Supplementary Table 1. Mitochondrial metabolism gene list

## STAR ⍰ Methods

### Key resource table

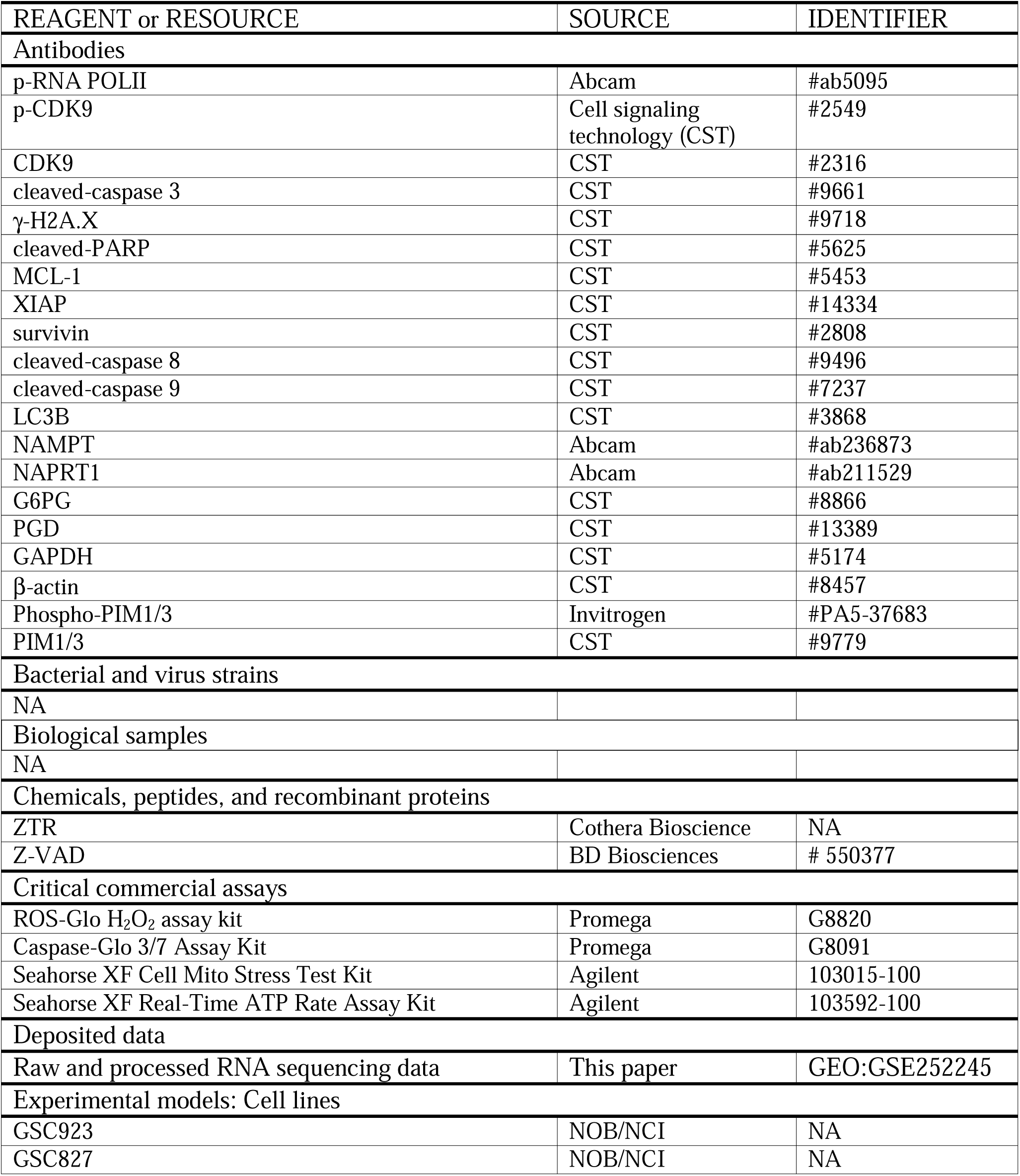
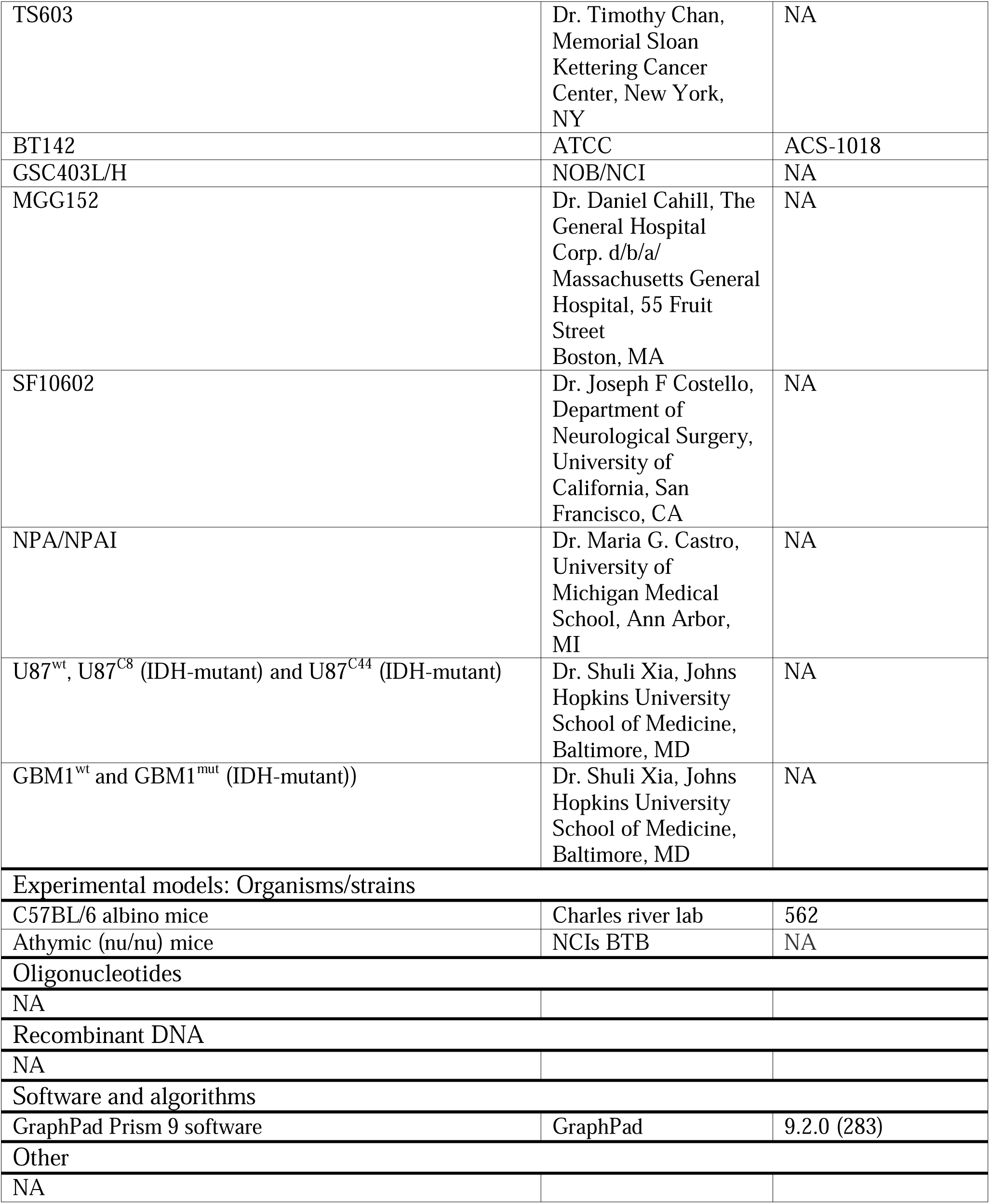

## RESOURCE AVAILABILITY

### Lead contact

Further information and requests for resources and reagents should be directed to and will be fulfilled by the Lead Contact, Jing Wu.

### Materials availability

Samples are available upon request, as collaboration, subject to availability.

### Data and code availability

Sequence data have been deposited at Gene Expression Omnibus (GEO) and are publicly available as of the date of publication (accession numbers GEO: GSE252245).

## EXPERIMENTAL METHODS DETAILS

### Cell culture and reagents

GSC923 and GSC827 were generated from patients’ glioblastoma tissues and cultured as previously reported ^5^. TS603 was obtained from Dr. Timothy Chan, Memorial Sloan Kettering Cancer Center, New York, NY ^46^. BT142 was purchased from ATCC and cultured as per instructions. NPA (shp53/shATRX/wtIDH1) and NPAI (shp53/shATRX/mIDH1^R132H^), mice-derived neurosphere cell lines, were provided by Dr. Maria G. Castro, University of Michigan Medical School, Ann Arbor, MI ^12^. Isogenic U87 (U87^wt^, U87^C8^ (IDH-mutant) and U87^C44^ (IDH-mutant)) and GBM1 (GBM1^wt^ and GBM1^mut^ (IDH-mutant)) cell lines were provided by Dr. Shuli Xia, Johns Hopkins University School of Medicine, Baltimore, MD ^47^. Briefly, the glioma stem-like cells were cultured in neurobasal-A medium or DMEM/F12 medium supplemented with bFGF, EGF, N2, B27, and penicillin/streptomycin at 37°C with 5% CO_2_. Adherent cell lines were cultured in DMEM medium supplemented with 10% FBS and 1% Penicillin/Streptomycin as previously reported ^5^.

### Kinase inhibition assay

In vitro kinase panel profiling was performed using the “HotSpot” assay platform by Reaction Biology Corp as described previously ^48^.

Briefly, kinase, substrate, cofactors were prepared in reaction buffer at room temperature. ZTR at 500nM, ATP and 33P ATP was added to a final concentration of 10 μM. Reaction spotting was performed using ion exchange filter paper. Any unbound phosphate was eliminated with phosphoric acid. The kinase activity results were reported as the percentage of remaining kinase activity in ZTR samples relative to the vehicle control.

### Cell viability assay

A total of 1 × 10^5^ cells/well were plated in geltrex precoated 12-well plates and treated with ZTR for 72 hours prior to cell counting using a Beckman Coulter ViCELL TM XR cell viability analyzer or Celigo Imaging Cytometer (Nexcelom Bioscience).

### High-throughput single agent screening

Six patient-derived glioma stem-like cell lines with *IDH1* mutations (GSC403L, GSC403H, BT142, MGG152, SF10602, and TS603) were screened with a cytotoxicity assay (48 hours Cell Titer-Glo) versus the MIPE 5.0 library (2481 agents: approved and investigational drugs with known MOA), as described and reported previously ^13,49^. Area under the curve (AUC) was utilized for ranking drug potency. For each compound tested, AUC values were averaged across the six cell lines and then normalized to Z-scores. MPO scores were plotted against Z-scores. Compounds of interest were highlighted based on an MPO score greater than 4 and an AUC Z-score of less than −3.

### ROS measurement

ROS was measured using ROS-Glo H_2_O_2_ assay kit (Promega). Briefly, 1 ×10^4^ cells/well were plated in geltrex pre-coated 96-well plates and treated with ZTR. H_2_O_2_ substrate solution was added in the culture medium and incubated for 6 hours. After adding ROS-Glo detection solution and incubating for 20 minutes, luminescence signal was read by the POLARStar Optima plate reader. The raw value was normalized to the protein content.

### Caspase 3/7 activity assay

Caspase 3/7 cleavage activity was measured by Caspase-Glo 3/7 Assay Kit (Promega). Cells were plated in a pre-coated 96-well plate. After treatment with ZTR for 72 hours, all wells were incubated with an equal volume of Caspase-Glo 3/7 reagent for 30 minutes and the luminescence signal was read by POLARStar Optima plate reader.

### Cell apoptosis assay

The ZTR-treated cells were differentially stained with APC Annexin V (BD Biosciences) and propidium iodide (PI) followed by flow cytometry analysis performed based on the manufacturer’s protocol. The fluorescence signal was measured by an BD LSRFortessa SORP I flow cytometer within 1 hour of staining and the percentage of apoptotic cells was quantified by FlowJo.

### Western blot

Cells were treated with ZTR at 15 nM prior to the cell harvest. The cell lysates were processed with ice-cold RIPA lysis buffer supplemented with protease and phosphatase Inhibitor Cocktail (Thermo Fisher Scientific). Protein concentration was determined by a DC Protein Assay kit (Bio-Rad) and the same amount of protein was resolved by NuPAGE 4%-12% Noves Bis-Tris gels (Invitrogen). Proteins were transferred to nitrocellulose membranes (Bio-Rad) and probed with primary antibodies (diluted 1:1000) at 4°C overnight. After secondary antibody incubation and washing, membranes were visualized using ChemiDoc imaging system (Bio-Rad).

### DNA fragmentation assay

Two million cells were treated with DMSO or 15nM of ZTR for 48 hours. DNA was extracted using the DNeasy Blood & Tissue kit (Qiagen) and total DNA concentration was determined by NanoDrop 8-Sample Spectrophotometer. Five hundred nanogram of DNA samples were resolved by electrophoresis in 10% Novex TBE gel (Invitrogen). The gel was stained by SYBR Gold DNA dye for 20 minutes and visualized by the ChemiDoc Imaging system (Bio-Rad).

### MitoTracker Green FM staining

Cells were seeded in pre-coated 8-well glass coverslip (BD Biosciences) and treated with DMSO or 15nM ZTR for 48 hours. After removing media from the coverslip, prewarmed staining solution with MitoTracker Green FM (100 nM, Thermo Fisher Scientific) was added for 30 minutes incubation at 37°C. After staining nuclei with Hoechst 33342, cells were observed using a NIKON confocal microscope.

### Blue native polyacrylamide gel electrophoresis and complex I in-gel activity assay

Cells were collected, washed with PBS, and homogenized in the sample buffer, supplemented with 1% digitonin. After centrifuging the cell lysate at 20,000 g for 30 min at 4°C, supernatant was collected and aliquoted for further assay. Protein concentration was measured by the DC Protein Assay kit (Bio-Rad) and the same amount of sample was mixed with sample buffer, 5% G-250 Sample Additive, and deionized water for native gel electrophoresis in dark blue cathode buffer. When gel electrophoresis was done, the gel was fixed in 40% methanol, 10% acetic acid buffer, and then the gel was washed in destain buffer (8% acetic acid) until the background was washed off for visualization by ChemiDoc Imaging system (Bio-Rad).

For complex I in-gel activity assay, after gel electrophoresis in light blue cathode buffer, the gel was incubated in fresh complex I substrate buffer (2 mM Tris·Cl, 0.1 mg/ml NADH, 2.5 mg/ml Nitrotetrazolium Blue chloride) for 20 min ^50^. The reaction was stopped by 10% acetic acid and the gel was washed with water for visualization by ChemiDoc Imaging system (Bio-Rad).

### Mitochondrial respiration and ATP real-time measurement assay

Mitochondria respiration and real-time ATP production was assessed by Seahorse XF Cell Mito Stress Test Kit (Agilent) and Seahorse XF Real-Time ATP Rate Assay Kit (Agilent), respectively. Briefly, cells were plated at a density of 40,000 cells/well in the pre-coated Seahorse cell culture plates. After cells were attached and stabilized, cells were treated with DMSO or ZTR for 24 hours. Medium in all wells was replaced by the XF based assay medium supplemented with 10mM glucose, 10mM sodium pyruvate and 2mM glutamine. After loading to the Seahorse XF96 analyzer (Agilent), compounds included in the Mito Stress Test Kit (oligomycin, FCCP and Rot/AA) or Real-Time ATP Rate Assay Kit (oligomycin and Rot/AA) were injected according to the pre-set programs for measurement.

### Transmission electron microscope (TEM) assay

GSC923 and TS603 cells were treated with DMSO or 15nM ZTR for 48 hours treatment for further TEM assay as previously described ^51^. Briefly, each sample in an individual well in the form of adherent cells was pre-fixed prior to be processed for TEM analysis. All samples were stained with 1% Osmium Tetroxide and 0.5% Uranyl Acetate successively. And then all samples were dehydrated and kept in 100% pure resin for infiltration overnight. Samples would be polymerized, and ultra-thin sectioned with Leica UC6 Microtome with 70-80 nm thickness and picked up on 150-Cu mesh grid. Following post-staining with 50:50 (0.5% Uranyl Acetate:70% Ethanol), samples would be carbon coated and then imaged by Hitachi TEM microscope H7600, with bottom mount camera at HV: 80.0kv for TEM analysis.

### RNA sequencing

GSC923 and TS603 cells were treated with DMSO or 15nM ZTR and collected after 24 or 48 hours. Total RNA was extracted using RNeasy Mini Kit (Qiagen) and passed the quality control of the Agilent RNA ScreenTape system.

mRNA-Seq samples were pooled and sequenced on NextSeq 2000 P3 using TruSeq Stranded mRNA Kit and paired-end sequencing. Reads were aligned to the reference genome hg38 and gene expression levels were quantified using STAR and RSEM tools. Raw RNA-Seq data filtered via the filter by expression function from the Edger package ^52^ was normalized with the default DESeq2 workflow ^53^. DESeq2 rlog transformed counts were filtered for protein coding genes only. Enrichment analysis was performed using Gene Set Enrichment Analysis (GSEA) on the filtered rlog counts with the Signal2Noise ranking metric ^54^. The Hallmark, Reactome, Kegg, and Gene Ontology biological process collections within the Molecular Signatures Database (MSigDB) were analyzed for enrichment with the GSEA algorithm ^55^. Only enriched gene sets with an FDR value less than .05 were considered for further analysis.

### Syngeneic and xenograft orthotopic murine models of glioma

All the animal experiments were reviewed and approved by the National Cancer Institute (NCI) Animal Use and Care Committee. Both syngeneic and xenograft orthotopic glioma mouse models were created to evaluate the treatment effect of ZTR in IDH-mutant and IDH-wt gliomas. To generate syngeneic model, isogenic cell line NPA (IDH-wt) or NPAI (IDH-mut) were injected (50,000 cells/2 μL) into the striatum of 8-weeks old C57BL/6 albino mice using a stereotactic device (coordinates, 2mm anterior and 2mm lateral from bregma, and 2.5 mm depth from the dura). Animals bearing IDH-mut and IDH-wt tumors were randomized into two groups based on body weights (n=7 mice for each group). Seven days after tumor implantation, ZTR (30mg/kg, i.p.) was administered twice a week for 21 days. To generate xenograft orthotopic model of gliomas with and without *IDH* mutation, isogenic cell line GBM1-wt or GBM1-mutant (500,000 cells/2 μl) were injected into 7-week-old female athymic (nu/nu) mice as previously described ^56^. Two weeks after injection, ZTR treatment (30mg/kg, i.p.) was administered, twice a week for 10 weeks. Animal behavior and body weight were monitored constantly. Animals were euthanized when they reached the endpoints. Tumor tissues were collected to analyze PD markers after treatment. Kaplan-Meier analysis was performed to determine the survival curve.

### Statistical analysis

Statistical analysis was performed using GraphPad Prism software using Student’s *t*-Test or one-way ANOVA test followed by Student’s t-Test as the post-statistical analysis. All tests were two-sided, and the results were shown as mean ± SEM or SD. For time course data, a two-sample Student’s t-test at each time point with the Bonferroni correction was performed. A p-value less than 0.05 was considered statistically significant. *, P < 0.05; **, P < 0.01; ***, or P < 0.001.

---

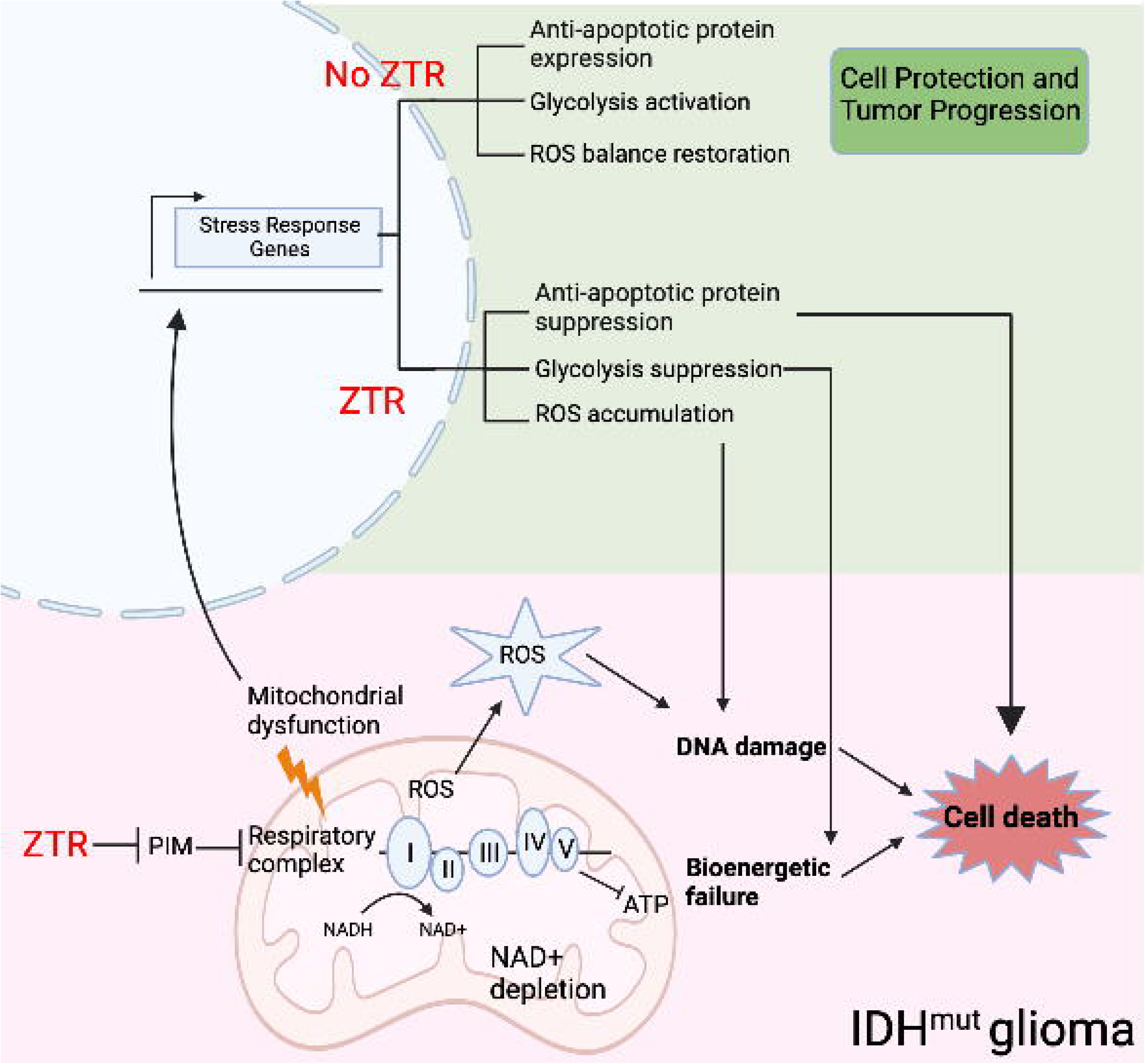

