## Supplementary figures and images for "Exploiting the therapeutic vulnerability of IDH-mutant gliomas with zotiraciclib"

### Supplementary figure 1

Suppl Figure 1

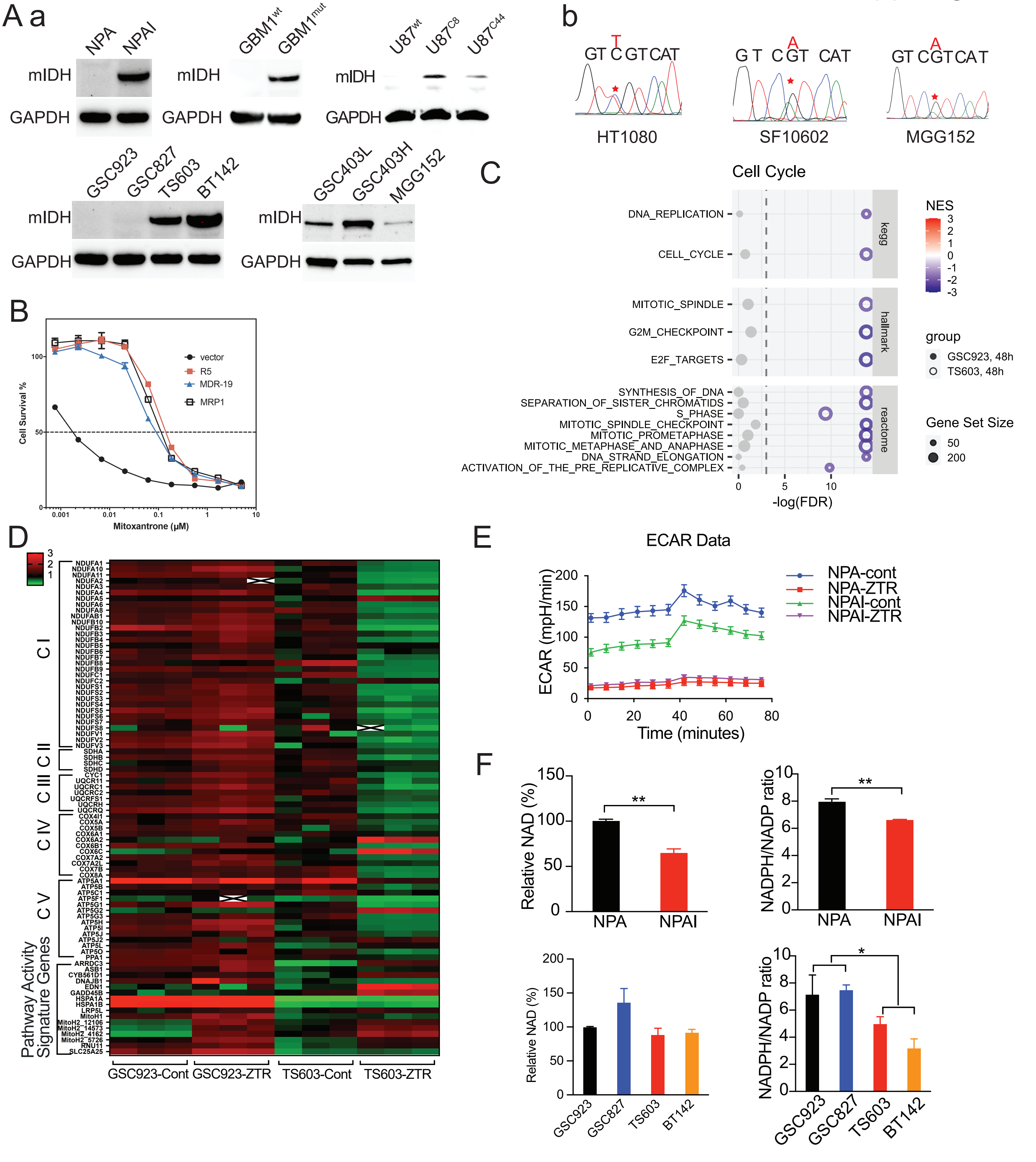

### Supplementary figure 2

Suppl Figure 2

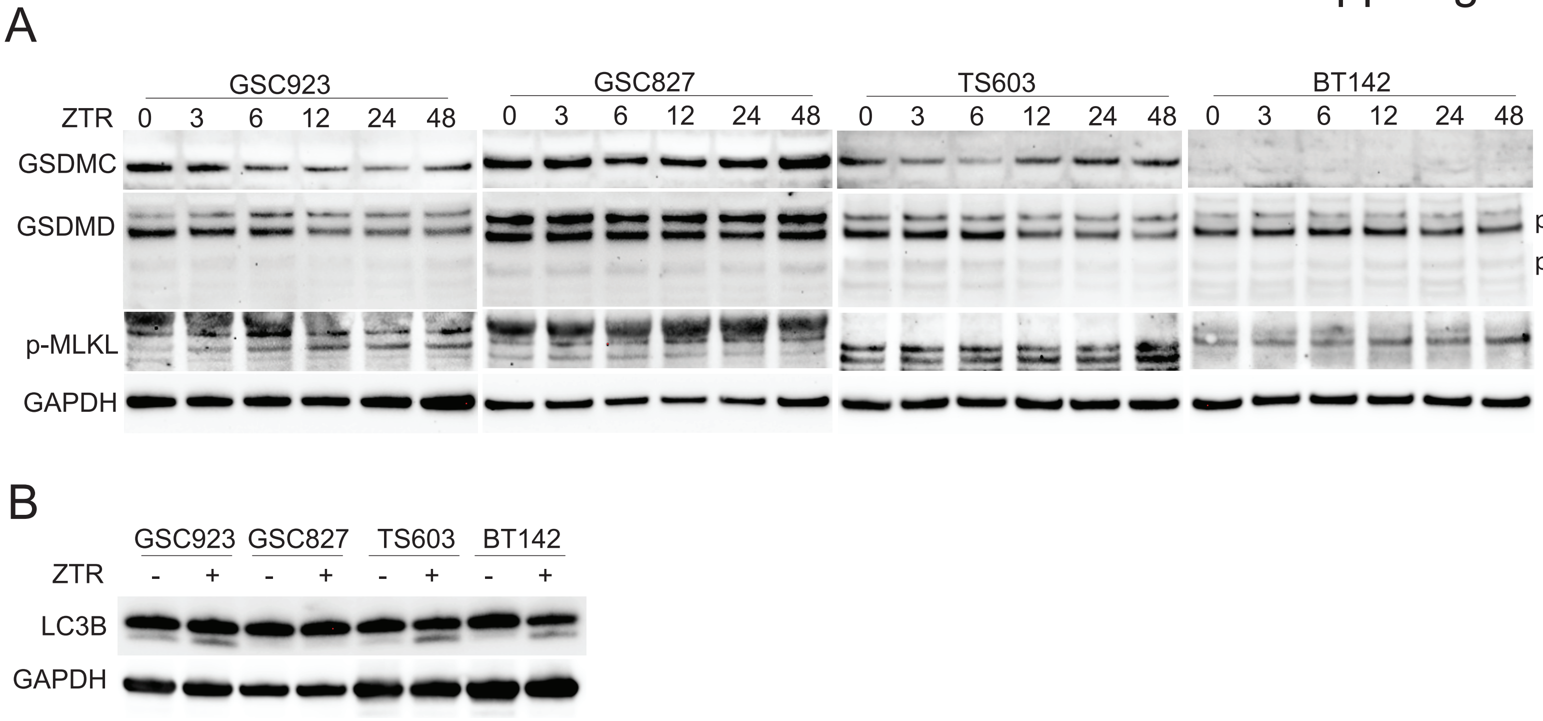
